# LSD1 inhibitors induce neuronal differentiation of Merkel cell carcinoma by disrupting the LSD1-CoREST complex and activating TGFβ signaling

**DOI:** 10.1101/2020.04.14.041657

**Authors:** Lukas Leiendecker, Pauline S. Jung, Tobias Neumann, Thomas Wiesner, Anna C. Obenauf

## Abstract

Merkel cell carcinoma (MCC) is a highly aggressive, neuroendocrine skin cancer that is either associated with the clonal integration of the Merkel cell polyomavirus or with chronic sun exposure^1,2^. Immunotherapy is initially effective in many patients with metastatic MCC, but the response is rarely durable^3,4^. MCC lacks actionable mutations that could be utilized for targeted therapies, but epigenetic regulators, which govern cell fate, provide unexplored therapeutic entry points. Here, we performed a pharmacological screen in MCC cells, targeting epigenetic regulators. We discovered that the lysine-specific histone demethylase 1A (LSD1/KDM1A) is required for MCC growth *in vitro* and *in vivo*. HMG20B (BRAF35), a poorly characterized subunit of the LSD1-CoREST complex, is also essential for MCC proliferation. LSD1 inhibition in MCC disrupts the LSD1-CoREST complex, directly induces the expression of key regulators of the neuronal lineage and of members of the TGFβ pathway, and activates a gene expression signature corresponding to normal Merkel cells. Our results provide a rationale for evaluating LSD1 inhibitors, which are currently being tested in patients with leukemia and solid tumors, in MCC.

## Main

Epigenetic regulators that maintain aberrant cell states have emerged as accessible entry points for targeted therapies^5,6^. To assess the therapeutic potential of epigenetic regulators in MCCs, we performed a pharmacological screen with 43 compounds targeting epigenetic modifiers in the MCC cell lines PeTa, WaGa, MKL-1, and MS-1 and human dermal fibroblasts (HDFB) (Fig. 1a and Supplementary Table 1). Compounds targeting EP300/CBP, BRD1/TAF1, and BET family proteins^7^ reduced MCC growth, but also reduced the growth of HDFBs, indicating a low specificity. In contrast, treatment with the LSD1 inhibitor (LSD1i) GSK-LSD1 strongly reduced the growth of the MCC cell lines, but not of HDFBs (**Fig. 1a and Supplementary Table 2**). We also treated cells with a structurally related LSD1i ORY-1001^8^ and observed similar, specific inhibition of cell growth (**Fig. 1b and Extended Data Fig. 1a**). The IC50 values for inhibition of MCC cell growth were in the low nM range for both GSK-LSD1 and ORY-1001, indicating that MCC cell lines have a high sensitivity to LSD1i (**Fig. 1c**).

**Fig. 1.**
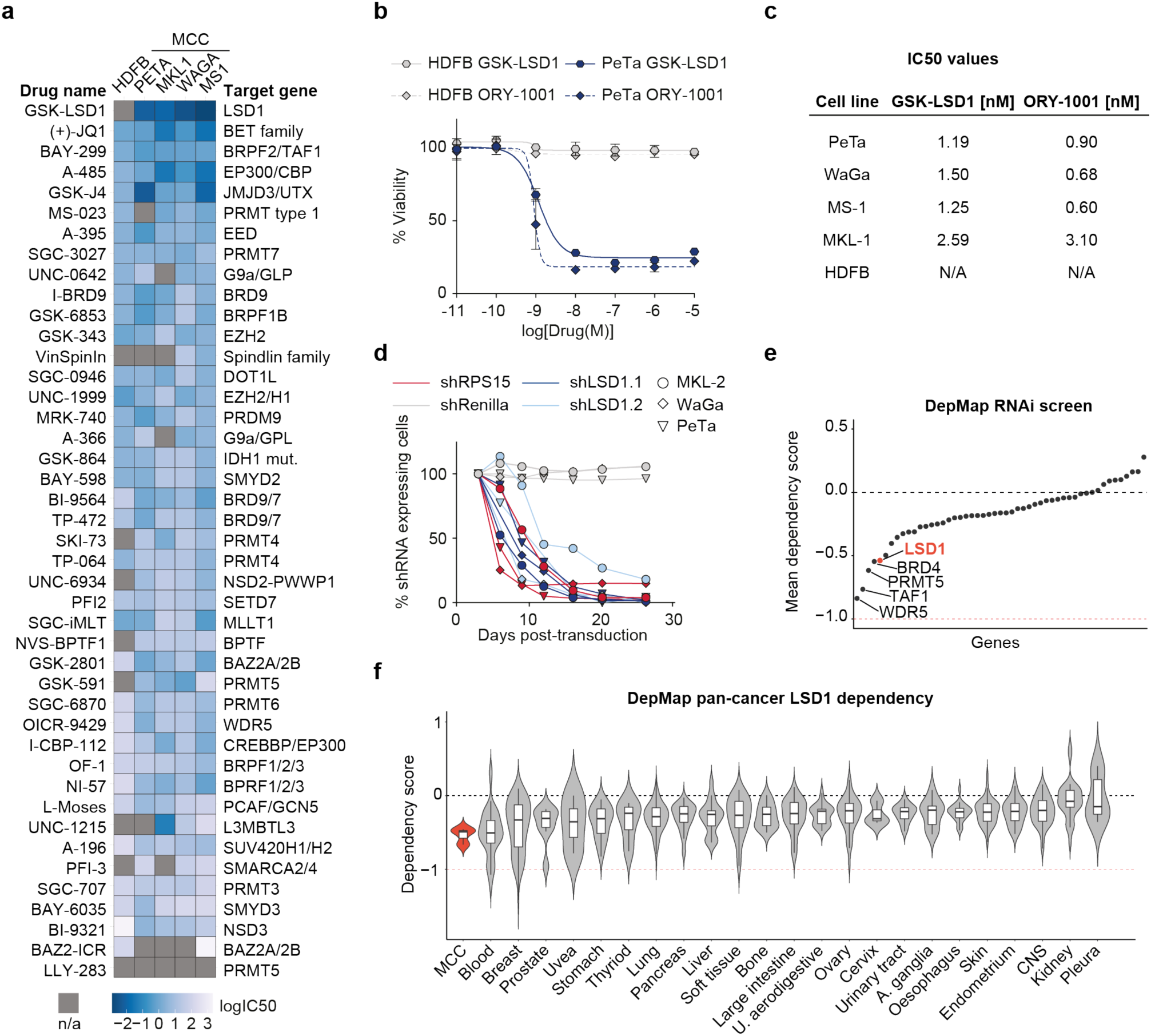
LSD1 is required for Merkel cell carcinoma proliferation. **a**, Heatmap of IC50 values for cell viability. Human dermal fibroblasts (HDFB) controls and 4 MCC cell lines (PeTa, MKL-1, WaGa, MS-1) were treated with the 43 indicated small molecules targeting epigenetic modifiers. n/a, IC50 values could not be calculated. **b**, Dose-response curves of PeTa cells and control HDFB cells after 6 days of treatment with GSK-LSD1 and ORY-1001. Dose-response curves of 3 other MCC cell lines are displayed in Extended Data Figure 1a. n = 4 technical replicates. Data are represented as means ±SEM. **c**, Calculated IC50 values for reduced growth of PeTa, WaGa, MS-1, MKL-1 as well as HDFB controls. **d**, *In vitro* competition assay of the 3 MCC cell lines MKL-2, PeTa, and WaGa transduced with either shLSD1.1, shLSD1.2, shRenilla (negative control), or shRPS15 (positive control). Individual graphs are displayed in Extended Data Figure 1e. **e**, Dependency plot depicting the mean dependency of the 3 MCC cell lines PeTa, MKL-1, MKL-2 on the genes targeted by the compound library in Fig. 1a. A score of 0 indicates that a gene is not essential; correspondingly -1 is comparable to the median of all pan-essential genes. Data obtained from DepMap; dependencies for the individual cell lines are displayed in Extended Data Figure 1f. **f**, Violin plot depicting the LSD1 dependency score in MCC compared to cancer types from 23 tissues, ordered according to mean dependency score. Boxplots: median at central line, first and third quartiles at lower and upper hinge and minimum and maximum values at whiskers. Data obtained from DepMap RNAi screen. Blood, hematopoietic and lymphoid tissue; U. aerodigestive, upper aerodigestive tract; A. ganglia, autonomic ganglia; CNS, central nervous system.

**Extended Data Fig. 1.**
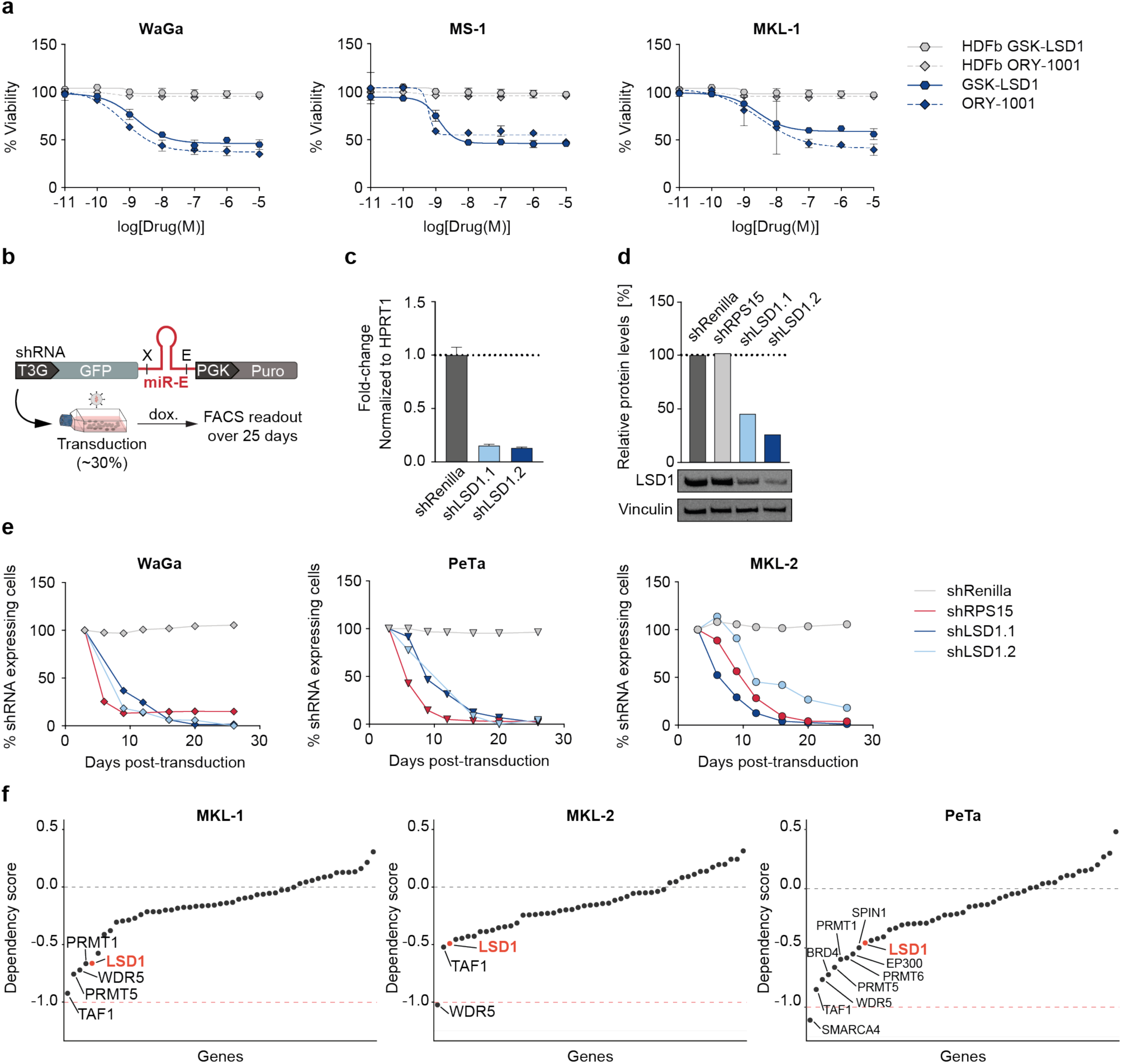
LSD1 is required for Merkel cell carcinoma proliferation. **a**, Dose-response curves for viability of 3 MCC cell lines (WaGa, MS-1, MKL-1) and control HDFB cells after 6 days of treatment with GSK-LSD1 and ORY-1001. n = 4 technical replicates for each sample. Data are represented as means ± SEM. **b**, Schematic depicting the *in vitro* shRNA-based competition assay. **c**, RT-qPCR of LSD1 RNA in the indicated shRNA-knockdown MKL-2 cells. n = 4 technical replicates. Bar graphs represent mean ± SD. **d**, Protein quantification of LSD1 protein levels and Western blot of LSD1 protein levels in MKL-2 cells. **e**, Individual graphs of the *in vitro* competition assay shown in Fig. 1d. The 3 MCC cell lines WaGa, PeTa, and MKL-2 were transduced with either shLSD1.1, shLSD1.2, shRenilla or shRPS15. **f**, Individual dependency plots of the 3 MCC cell lines PeTa, MKL-1, MKL-2 for the genes targeted by the compound library in Fig. 1a. A score of 0 indicates that a gene is not essential; correspondingly -1 is comparable to the median of all pan-essential genes. Data obtained from DepMap.

To verify that the effects of LSD1i treatment reflect specific inhibition of LSD1, we used RNAi to deplete LSD1 in MCCs. Briefly, the MCC lines were engineered to express doxycycline-inducible shRNAs targeting LSD1 (shLSD1.1 or shLSD1.2), Renilla luciferase (negative control) or the ribosomal protein RPS15 (positive control) (**Extended Data Fig. 1b**). The LSD1-targeting shRNAs, but not the control shRNAs, reduced LSD1 RNA levels to ∼15% and protein levels to ∼30% (**Extended Data Fig. 1c,d**). Importantly, the LSD1-targeting shRNAs and shRPS15 strongly reduced cell growth in all the tested MCC cell lines (**Fig. 1d and Extended Data Fig. 1e**), whereas shRenilla had no effect. Additionally, we analyzed independent, genome-wide RNAi screening data from the DepMap project^9^, which includes genetic vulnerability maps for the MCC cell lines PeTa, MKL-1, and MKL-2. We examined the genes encoding the epigenetic regulators from our initial screen and found that LSD1 scores among the top 5 dependencies for MCC proliferation (**Fig. 1e and Extended Data Fig. 1f**). To assess the specificity of LSD1 dependency, we compared MCC to other cancer types and found that the mean LSD1 dependency score of MCC was similar to that of hematopoietic and lymphoid malignancies (**Fig. 1f**). Intriguingly, subtypes of hematopoietic and lymphoid cancer are known to respond to LSD1i^10,11^. Collectively, our data indicate that genetic and pharmacological inhibition of LSD1 inhibits MCC cell growth.

To assess the therapeutic efficacy of LSD1i *in vivo*, we subcutaneously injected PeTa cells into the flanks of immunocompromised NSG mice. After the tumors reached a volume > 50mm^3^ (16 days post-injection), we treated the mice with GSK-LSD1 or vehicle (**Fig. 2a**). We found that subsequent tumor growth was substantially reduced in mice treated with LSD1i compared to vehicle (**Fig. 2b and Extended Data Fig. 2a**). Mouse weight remained relatively stable throughout the treatment, suggesting good tolerability (**Extended Data Fig. 2b**). The tumor control in all LSD1i treated mice resulted in 100% overall survival, whereas all vehicle-treated mice had to be euthanized due to high tumor burden (**Fig. 2c,d**).

**Fig. 2.**
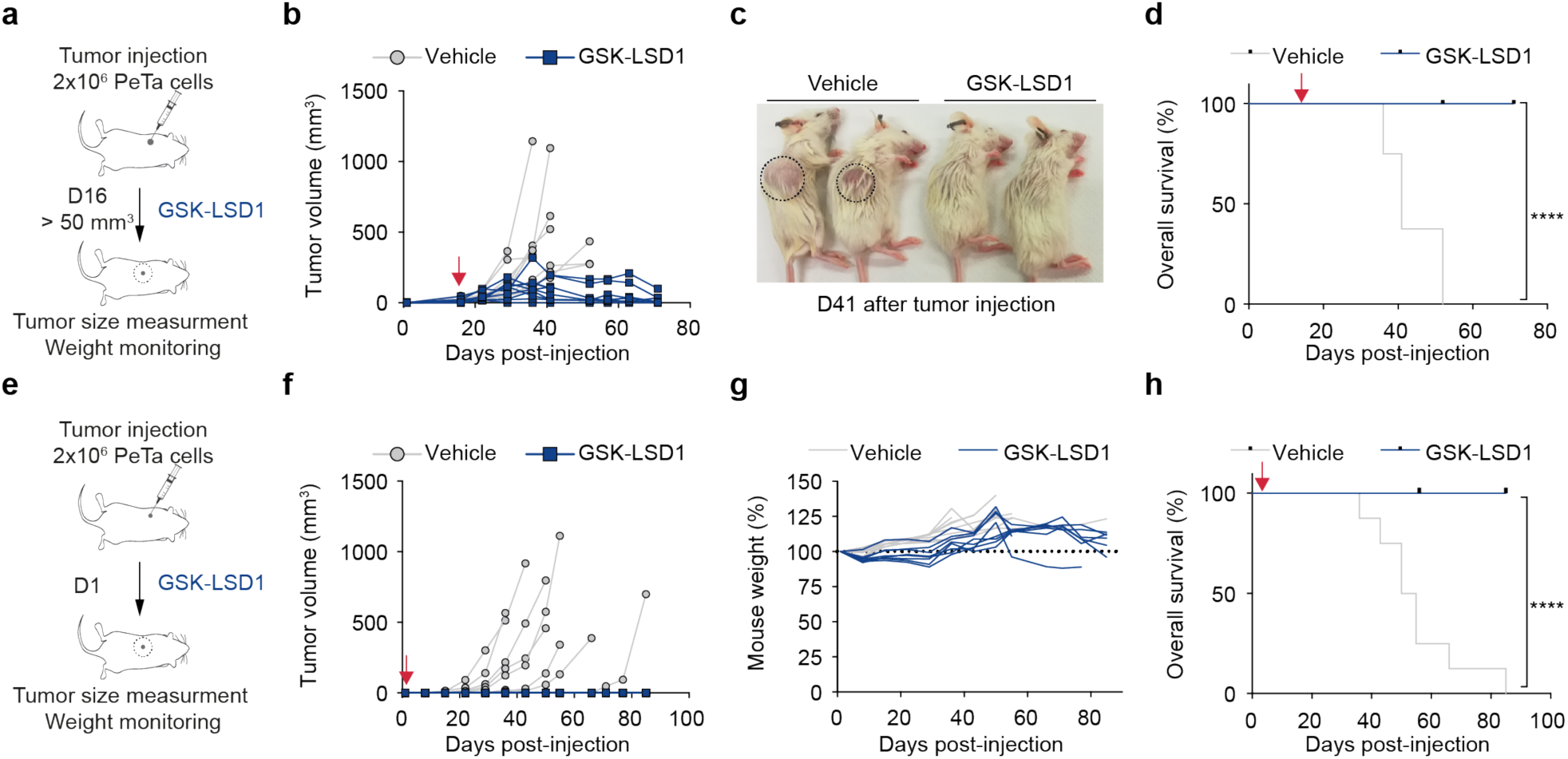
Pharmacological LSD1 inhibition controls tumor growth *in vivo*. **a**, Schematic depicting the experimental setup for *in vivo* xenograft treatment of MCC tumors with GSK-LSD1 in NSG mice. GSK-LSD1 or vehicle treatment was started 16 days after PeTa cell injection, when tumor volume was > 50mm^3^. **b**, Individual tumor growth in GSK-LSD1 (n=9) or vehicle-treated (n=8) mice. Red arrow: start of therapy, day 16. **c**, Representative picture of mice treated with vehicle or GSK-LSD1 at day 41 after PeTa cell injection. Tumor location is indicated with a circle. **d**, Kaplan-Meier curve. Mice were sacrificed when tumors reached a volume > 1.5cm^3^ or a dimension > 1.5cm. Red arrow: start of therapy, day 16. ****p < 0.0001 (log-rank Mantel-Cox test). **e**, Schematic depicting the experimental setup for *in vivo* xenograft treatment of MCC “micrometastases” with GSK-LSD1 in NSG mice. GSK-LSD1 or vehicle treatment was started one day after tumor injection (D1). **f**, Individual tumor growth in GSK-LSD1 (n=8) or vehicle-treated (n=8) mice. Red arrow: start of therapy, day 1. **g**, Relative mouse weight (%) during treatment. **h**, Kaplan-Meier curve. Mice were sacrificed when tumors reached a volume > 1.5cm^3^ or a dimension > 1.5cm. Red arrow: start of therapy, day 1. ****p < 0.0001 (log-rank Mantel-Cox test).

**Extended Data Fig. 2.**
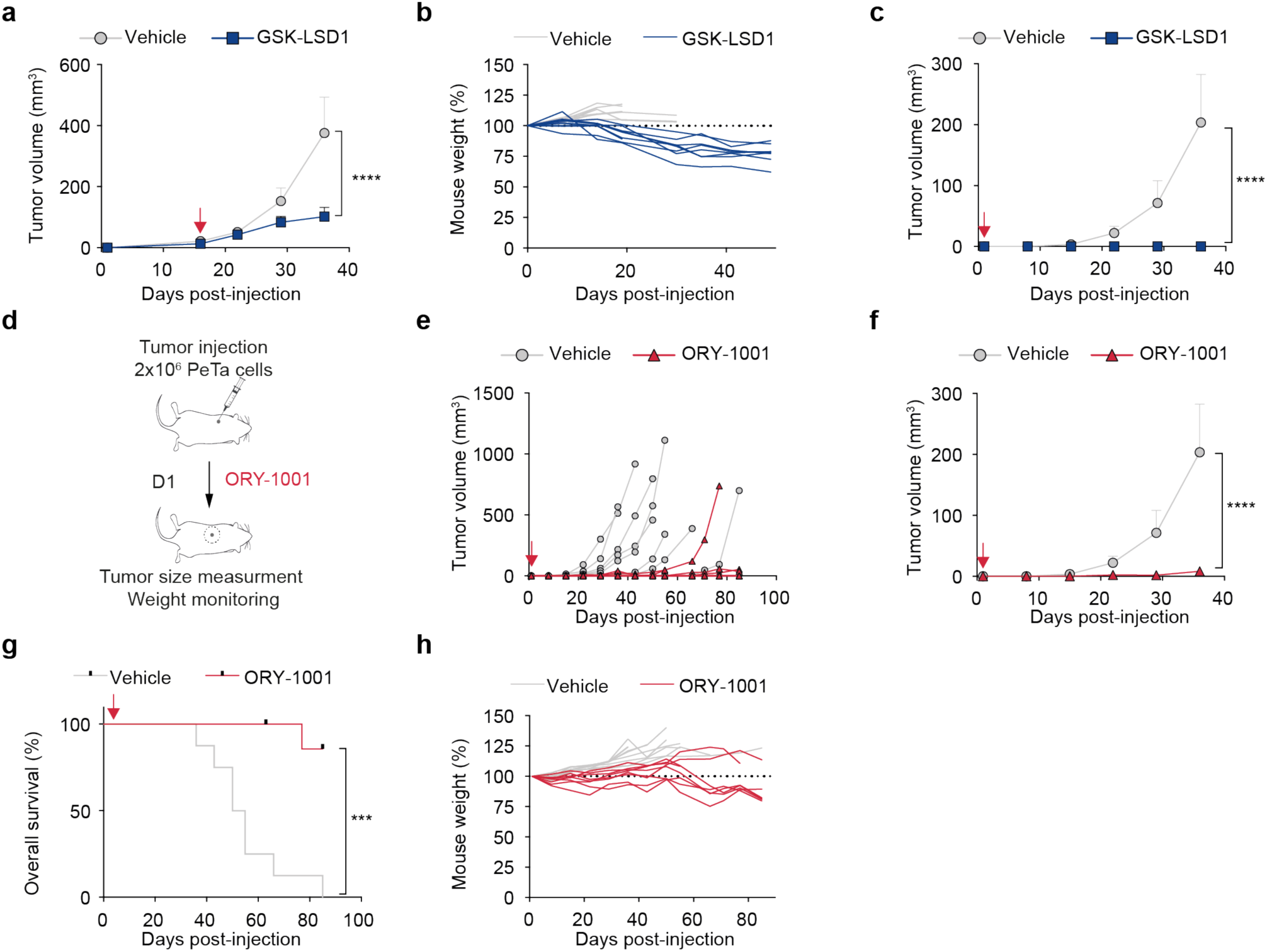
Pharmacological LSD1 inhibition controls tumor growth *in vivo*. **a**, Treatment response of subcutaneous tumors in mice treated with GSK-LSD1 (n=9) or vehicle (n=8); arrow, start of therapy, day 16. Data are represented as means ±SEM. ****p<0.0001, two-way ANOVA. **b**, Relative mouse weight (%) during treatment. **c**, Treatment response of subcutaneous tumors treated with GSK-LSD1 (n=8) or vehicle (n=8); Red arrow: start of therapy, day 1. Data are represented as means ±SEM. ****p<0.0001, two-way ANOVA. **d**, Schematic depicting the experimental setup for *in vivo* xenograft tumor treatment with ORY-1001 in NSG mice. ORY-1001 or vehicle treatment was started one day after tumor injection (D1). **e**, Individual tumor growth with ORY-1001 (n=8) or vehicle-treated (n=8) mice. Red arrow: start of therapy, day 1. **f**, Treatment response of subcutaneous tumors treated with ORY-1001 (n=8) or vehicle (n=8); Red arrow: start of therapy, day 1. Data are represented as means ±SEM. ****p<0.0001, two-way ANOVA. **g**, Kaplan-Meier curve. Mice were sacrificed when tumors reached a volume > 1.5cm^3^ or a dimension > 1.5cm. Red arrow: start of therapy, day 1. ****p < 0.0001 (log-rank Mantel-Cox test). **h**, Relative mouse weight (%) during treatment.

MCC is a highly metastatic cancer involving all organs, which contributes to its high morbidity and mortality^12^. Once seeded, micro-metastases must grow and establish a tumor. To model the response of micro-metastases, we subcutaneously injected MCC cells and started LSD1i treatment one day post-injection, prior to tumor establishment (**Fig. 2e**). All mice treated with GSK-LSD1 (8/8) (**Fig. 2f-h and Extended Data Fig. 2c**), and most mice treated with ORY-1001 (7/8) (**Extended Data Fig. 2d-h**) remained tumor-free; in contrast, all vehicle-treated mice (8/8) grew tumors. Collectively, these data suggest LSD1 inhibition reduces MCC growth *in vitro* and *in vivo* and is a vulnerability in MCC that could be exploited therapeutically.

LSD1 acts in cell context-specific protein complexes to regulate gene expression by demethylating H3K4me1/2, H3K9me1/2, and non-histone proteins^13–15^. To uncover LSD1 binding partners in MCC, we immunoprecipitated LSD1, identified interacting proteins by mass spectrometry and performed unsupervised protein-protein interaction enrichment analysis (**Fig. 3a-c and Extended Data Fig. 3a**). We found that LSD1 interacts with members of the LSD1-CoREST corepressor complex (also called BRAF–histone deacetylase complex, BHC). In particular, we identified core members, including the histone deacetylase HDAC2, and RCOR1, RCOR2, RCOR3, which serve as a scaffold for complex assembly, as well as non-canonical members, including GSE1, HMG20A, HMG20B (F**ig. 3c and Extended Data Fig. 3a**). Importantly, we found that LSD1i treatment of MCC leads to dissociation of this complex, which is in line with studies in leukemias^16,17^ (**Fig. 3d,e**).

**Fig. 3.**
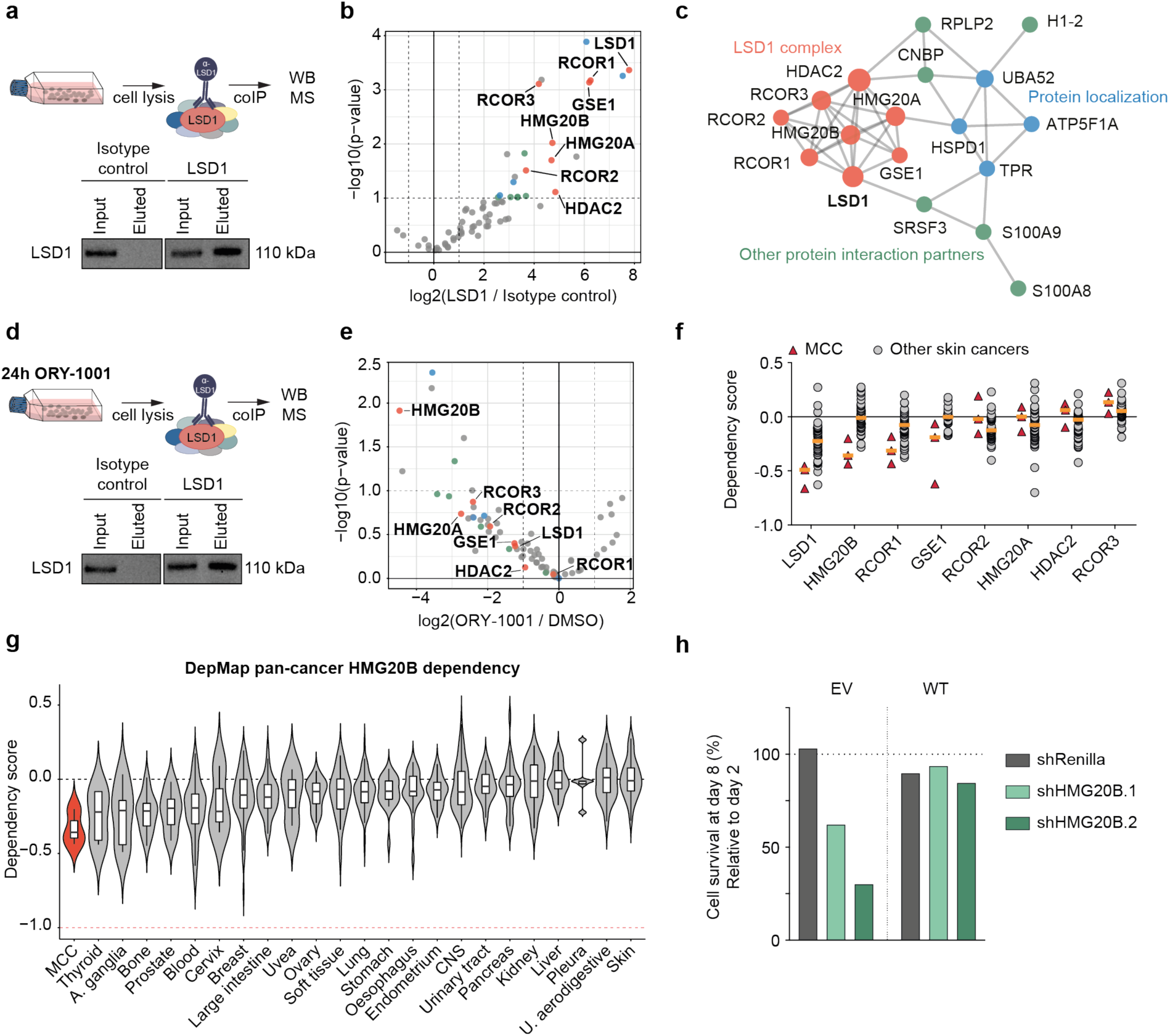
HMG20B is part of the LSD1-CoREST complex in MCC and is essential for proliferation. **a**, Schematics and Western blot of LSD1 co-immunoprecipitation (coIP) from PeTa cells. WB, Western blot; MS, mass spectrometry. **b**, Volcano plot displaying the protein-protein binding partners of LSD1. Proteins belonging to the LSD1 complex are marked in red, protein localization in blue and other protein interaction partners in green. **c**, Protein-protein interaction (PPI) mapping of the identified LSD1 binding partners. Individual complexes and p-values are displayed in Extended Data Fig. 3a. **d**, Schematics and Western blot of LSD1 co-immunoprecipitation (coIP) in ORY-1001 (1 μM) treated PeTa cells. WB, Western blot; MS, mass spectrometry. **e**, Volcano blot displaying the protein-protein binding partners of LSD1 depleting upon PeTa cells treated with ORY-1001 versus DMSO. Proteins belonging to the LSD1 complex are marked in red, protein localization in blue and other protein interaction partners in green. **f**, Dependency on LSD1 complex members in MCC compared to other skin cancers. Data obtained from DepMap. Median is indicated with a horizontal line. **g**, Violin plot depicting the HMG20B dependency in MCC compared to cancer types from 23 tissues, ordered according to mean. Boxplots: median at central line, first and third quartiles at lower and upper hinge and minimum and maximum values at whiskers. Data obtained from DepMap. Blood, hematopoietic and lymphoid tissue; U. aerodigestive, upper aerodigestive tract; A. ganglia, autonomic ganglia; CNS, central nervous system. **h**, Bar graph of HMG20B rescue experiment in PeTa cells transduced with either shRNAs targeting the endogenous HMG20B or Renilla (negative control) and with an overexpression construct expressing WT HMG20B. Cell survival is depicted at day 8 after transduction and relative to day 2. EV, empty vector control, WT, wild-type.

**Extended Data Fig. 3.**
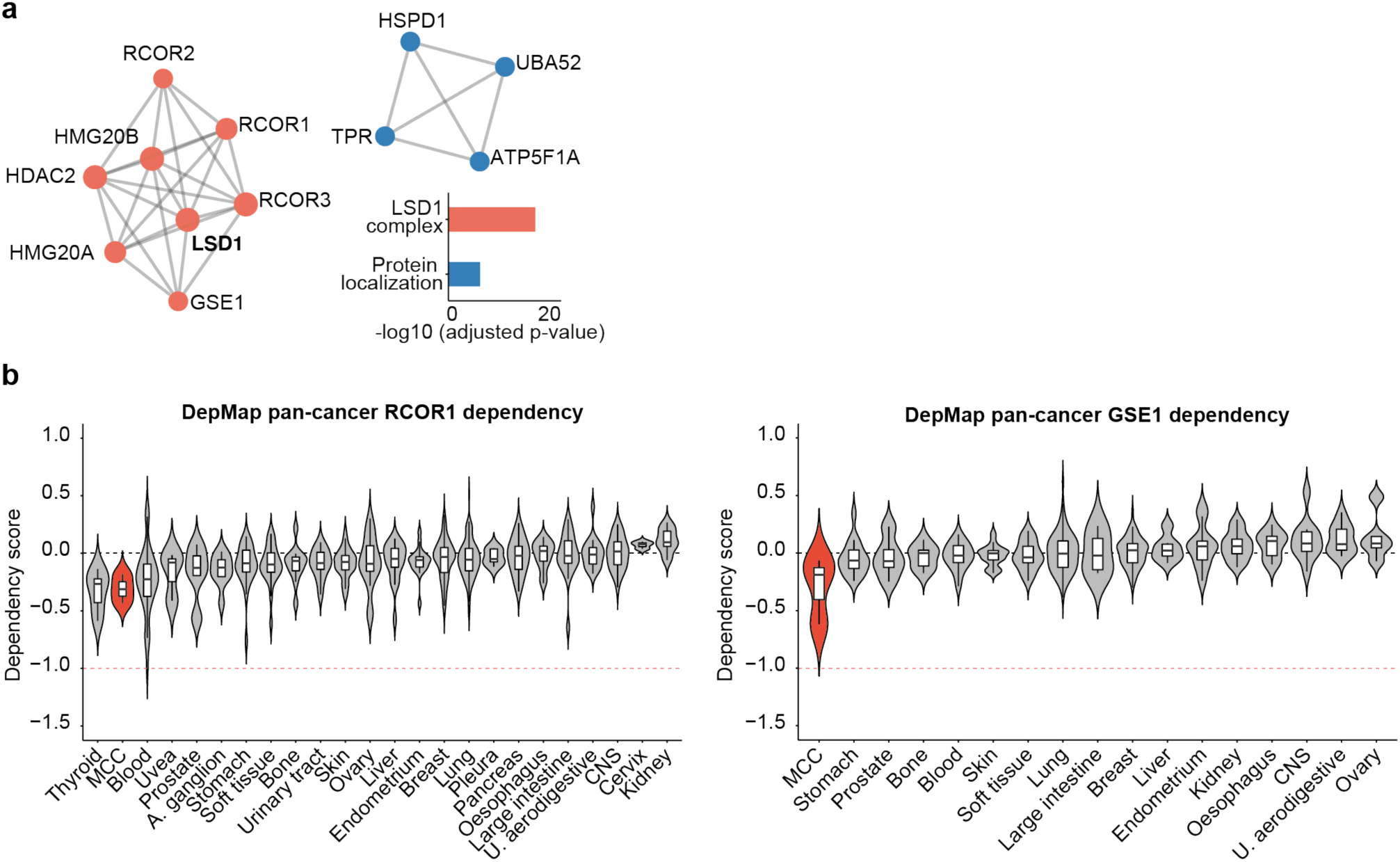
Cancer-type specific dependencies of the CoREST-complex members. **a**, Individual protein-protein interaction complexes identified by LSD1 co-immunoprecipitation/mass spectrometry and adjusted p-values. **b**, Violin plot depicting the dependency scores of RCOR1 (left) and GSE1 (right) in MCC compared to cancer types from 23 tissues, ordered according to mean dependency score. Boxplots: median at central line, first and third quartiles at lower and upper hinge and minimum and maximum values at whiskers. Data obtained from DepMap. Blood, hematopoietic and lymphoid tissue; U. aerodigestive, upper aerodigestive tract; A. ganglia, autonomic ganglia; CNS, central nervous system.

To determine if these LSD1 binding partners are also required for cell growth in MCC, we examined again the RNAi data from the DepMap project^9^. Our analysis suggested that LSD1, RCOR1, and HMG20B are specifically required in MCC compared to other cancers (**Fig. 3f,g and Extended Data Fig. 3b**). To verify that HMG20B is required for MCC growth, we depleted endogenous HMG20B. Indeed, depletion of HMG20B reduced MCC cell growth, which was rescued by expression of a WT HMG20B that is not targeted by the shRNA (**Fig. 3h**). Together, these data suggest that an LSD1-RCOR1-HMG20B complex is required for MCC growth.

When investigating how LSD1 controls MCC growth, we first noticed that MCC cells became smaller and formed dense clusters upon LSD1i treatment, as opposed to their typical growth as loose aggregates in suspension clusters (**Fig. 4a and Extended Data Fig. 4a**). However, we detected no evidence for apoptotic cell death, indicated by a lack of cleaved caspase-3/7 and PARP1 (**Fig. 4b and Extended Data Fig. 4b**). To investigate the molecular programs associated with LSD1i mediated growth inhibition, we performed RNAseq of DMSO- and GSK-LSD1-treated MCC cells after 6 days. We found that GSK-LSD1 treatment led to the upregulation of 870 genes, and the downregulation of 533 genes (**Fig. 4c**). We examined the promoter motifs in the deregulated genes and found that the Co-/Rest binding motif was enriched in the promoters of genes that become upregulated in MCCs upon inhibition of LSD1, indicating a direct regulation of these genes by the LSD1-CoREST complex (**Fig. 4d**). Pathway enrichment analysis of the upregulated genes was dominated by terms associated with nervous system development and neuronal differentiation programs (**Fig. 4e,f**). In contrast, cell cycle genes and the proliferation marker Ki-67 were downregulated (**Fig. 4g,h and Extended Data Fig. 4c,d**). Taken together, these data suggest that the therapeutic effect of LSD1i treatment is mediated by the expression of neuronal genes and differentiating MCC cells into a neuronal lineage. Interestingly, HMG20B, which we identified as a subunit of the LSD1-CoREST complex in MCC, was shown to be required to maintain full repression of neuron-specific genes^18,19^.

**Fig. 4.**
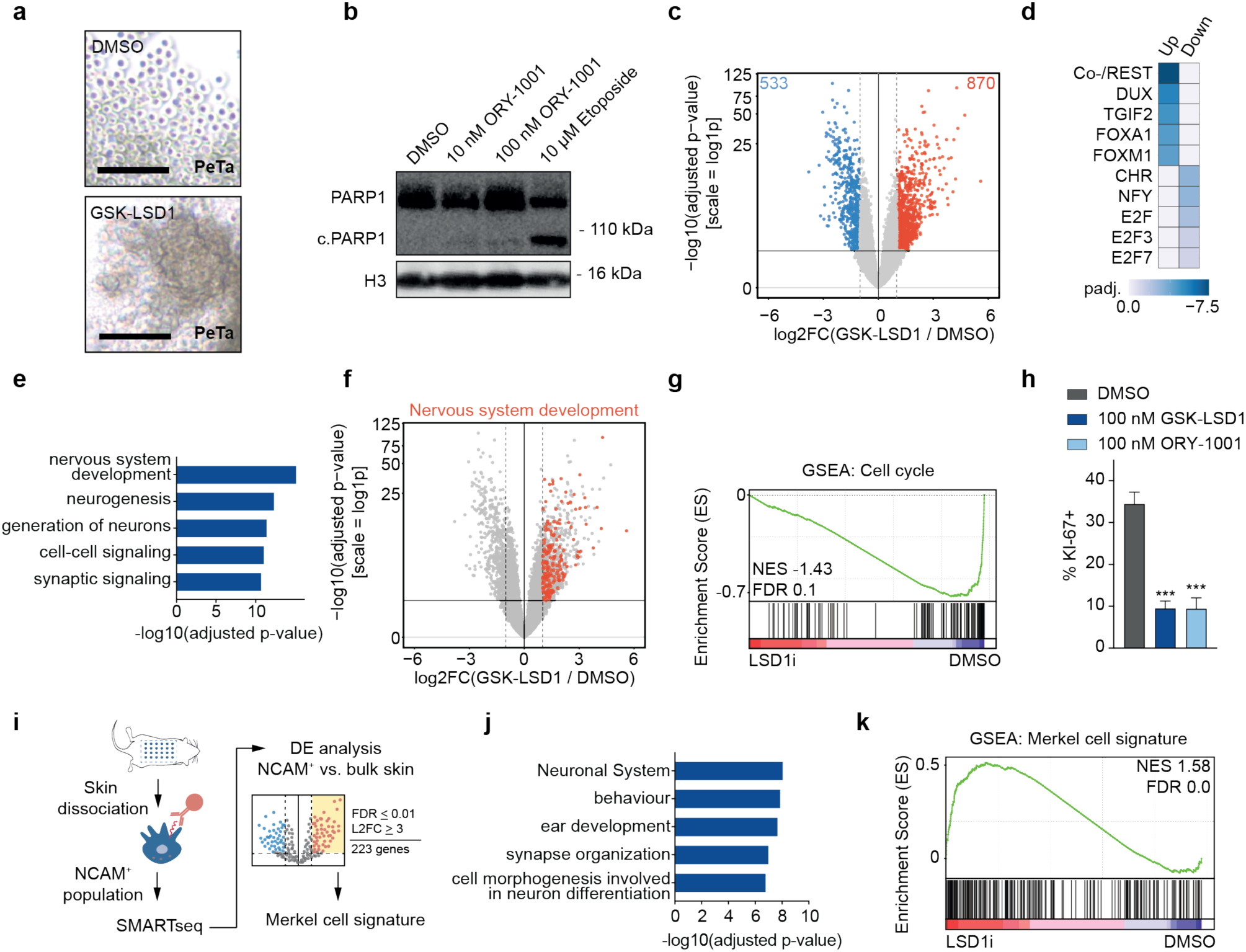
LSD1 inhibition de-represses neuronal differentiation genes. **a**, Photomicrographs of the MCC cell line PeTa after 6 days of GSK-LSD1 (100 nM) or DMSO treatment. Scale bar, 100 μm. **b**, Western blot of PARP1 cleavage. PeTa cells were treated for 6d with the indicated concentrations of ORY-1001. Etoposide serves as positive control for apoptosis, H3 serves as loading control. c.PARP1, cleaved PARP1. **c**, Volcano plot showing the -log10(adj.p-value) and log2fold-change (log2FC) for transcripts detected by RNAseq analysis of PeTa cells treated with 100 nM GSK-LSD1or DMSO after 6 days. Significantly up- and down-regulated genes (FDR<0.05; abs(log2FC)>1) and the corresponding number are marked in red and blue, respectively. **d**, Upstream gene promoter motif enrichment of up- and down-regulated genes upon GSK-LSD1 treatment displayed in Fig. 4c. padj, adjusted p-value; Up, upregulated genes; Down, downregulated genes. **e**, Pathway analysis of 870 genes significantly upregulated upon GSK-LSD1 displayed in Fig. 4c. **f**, Volcano plot displaying the upregulated and downregulated genes of Fig. 4c where genes involved in nervous system development (GO:0007399) are highlighted in red. **g**, Gene set enrichment analysis (GSEA) of cell cycle gene set of data in Fig. 4c. NES, normalized enrichment score; FDR, false discovery rate. **h**, KI-67 staining of PeTa cells treated for 3 days with GSK-LSD1 (1 μM) and ORY-1001 (1 μM). Data are represented as means ±SD. ***p < 0.001 (Student’s t-test). **i**, Schematic of Merkel cell extraction and Merkel cell signature generation. DE, differential expression; FDR, false discovery rate; L2FC, log2 fold-change. **j**, Pathway analysis of 223 genes upregulated in Merkel cell signature. **k**, Gene set enrichment analysis (GSEA) of the generated Merkel cell signature in the data in Fig. 4c. NES, normalized enrichment score; FDR, false discovery rate.

**Extended Data Fig. 4.**
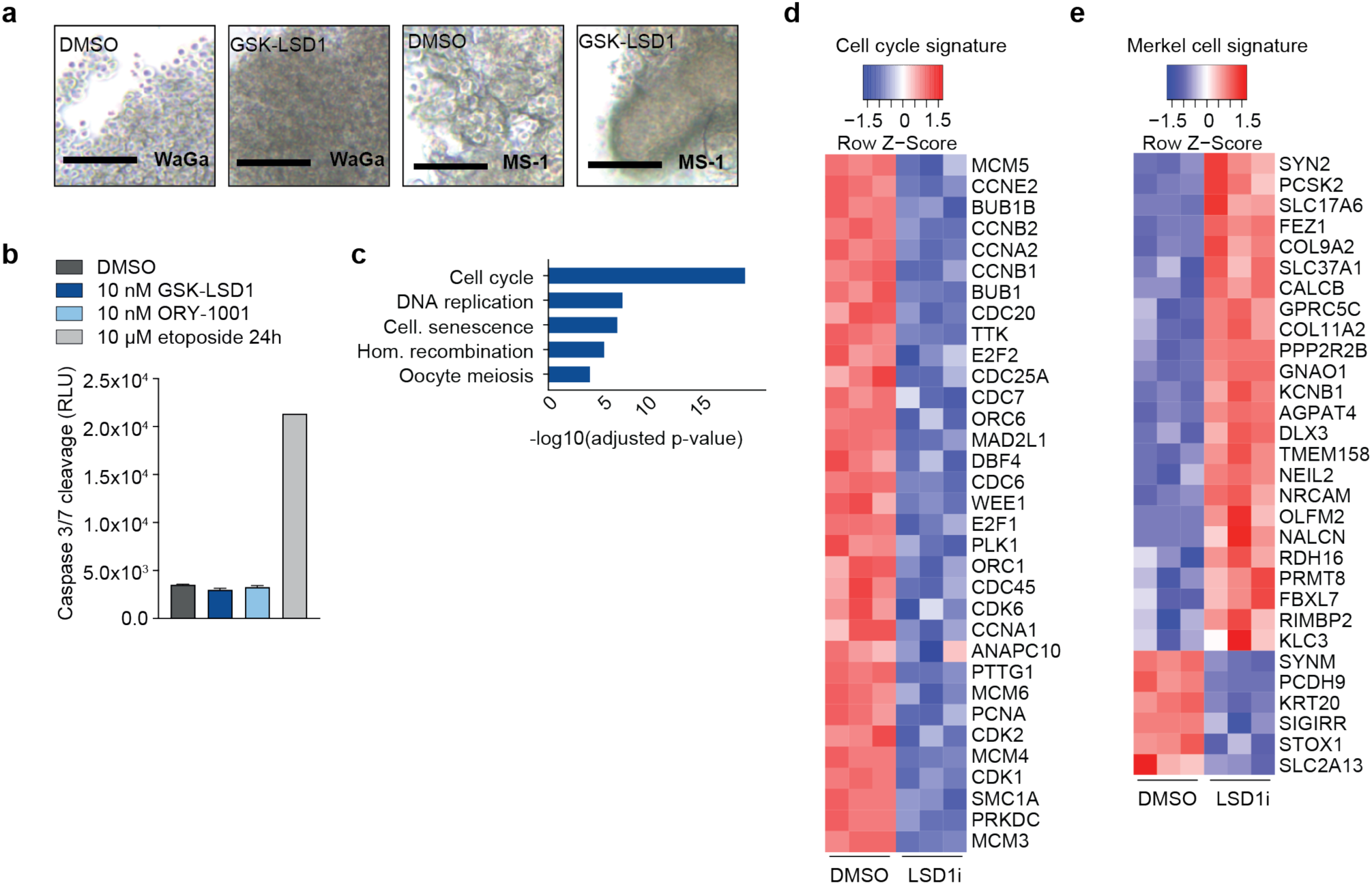
LSD1i down-regulate cell cycle genes and induce differentiation of MCC towards a Merkel cell fate. **a**, Photomicrographs of the MCC cell lines WaGa and MS-1 after 6 days of GSK-LSD1 (100 nM) or DMSO treatment. Scale bar, 100 μm. **b**, Caspase3/7 cleavage activity for PeTa cell line after 6 days of treatment with GSK-LSD1 or ORY-1001. Etoposide serves as a positive control for apoptosis. n = 3 technical replicates. Data are represented as means ±SD. RLU, relative luminescence units. **c**, Pathway analysis of 533 genes significantly downregulated upon 100 nM GSK-LSD1 for 6 days of data displayed in Fig. 4c. **d**, Heatmap of genes enriching in cell cycle pathway from the dataset in Fig. 4c. **e**, Heatmap of genes enriching in Merkel cell signature from the dataset in Fig. 4c.

Many genes of the neuronal lineage are expressed in normal Merkel cells, which are discussed to be a putative cell of origin of MCC^20^. To test whether LSD1i-treated MCC cells resemble normal Merkel cells, we performed low RNA-input transcriptome analysis (SMARTseq) to analyze the transcriptomes of Merkel cells purified from mouse skin and of bulk skin cells (**Fig. 4i**). Compared to the bulk skin transcriptome, the Merkel cell transcriptome was enriched for terms associated with neuronal differentiation and included genes important for normal Merkel cell development and maintenance, such as *ATOH1, SOX2*, and *INSM1* (**Fig. 4j and Supplementary Table 11**). We defined a “Merkel cell signature” comprising the top differentially expressed genes and applied it to the transcriptome of LSD1i-treated MCC (**Fig. 4k**). We found that the Merkel cell signature was highly enriched in LSD1i-treated MCC cells (**Fig. 4k and Extended Data Fig. 4e**), suggesting that LSD1i treatment induces differentiation of MCC towards a Merkel cell fate.

To interrogate the direct transcriptional responses to LSD1i treatment, we performed SLAMseq, which detects newly transcribed (nascent) RNA^21,22^. We treated MCC cells with vehicle or GSK-LSD1 for 30 min, added the uridine analog 4sU for 1 h or 6 h to label nascent transcripts, and performed SLAMseq (**Fig. 5a**). We observed a small set of genes with altered transcription after just 1 h, which likely represent direct targets of LSD1 in MCC cells (**Fig. 5b and Extended Data Fig. 4a,b**). Interestingly, both time points showed increased transcription of genes associated with neuron development (GO:0048666), including the transcription factors NEUROD1 and INSM1, which regulate neuronal and neuroendocrine differentiation, respectively, and are also expressed in normal Merkel cells (**Extended Data Fig. 5c and Supplementary Table 11**). In line with the role of LSD1 as a transcriptional repressor, the immediately upregulated LSD1 target genes remained upregulated throughout LSD1i treatment, whereas the initial downregulation of some LSD1 target genes was not maintained (**Fig. 5c**). Overall, these data suggest that LSD1i treatment directly upregulates the transcription of neuronal differentiation genes.

**Fig. 5.**
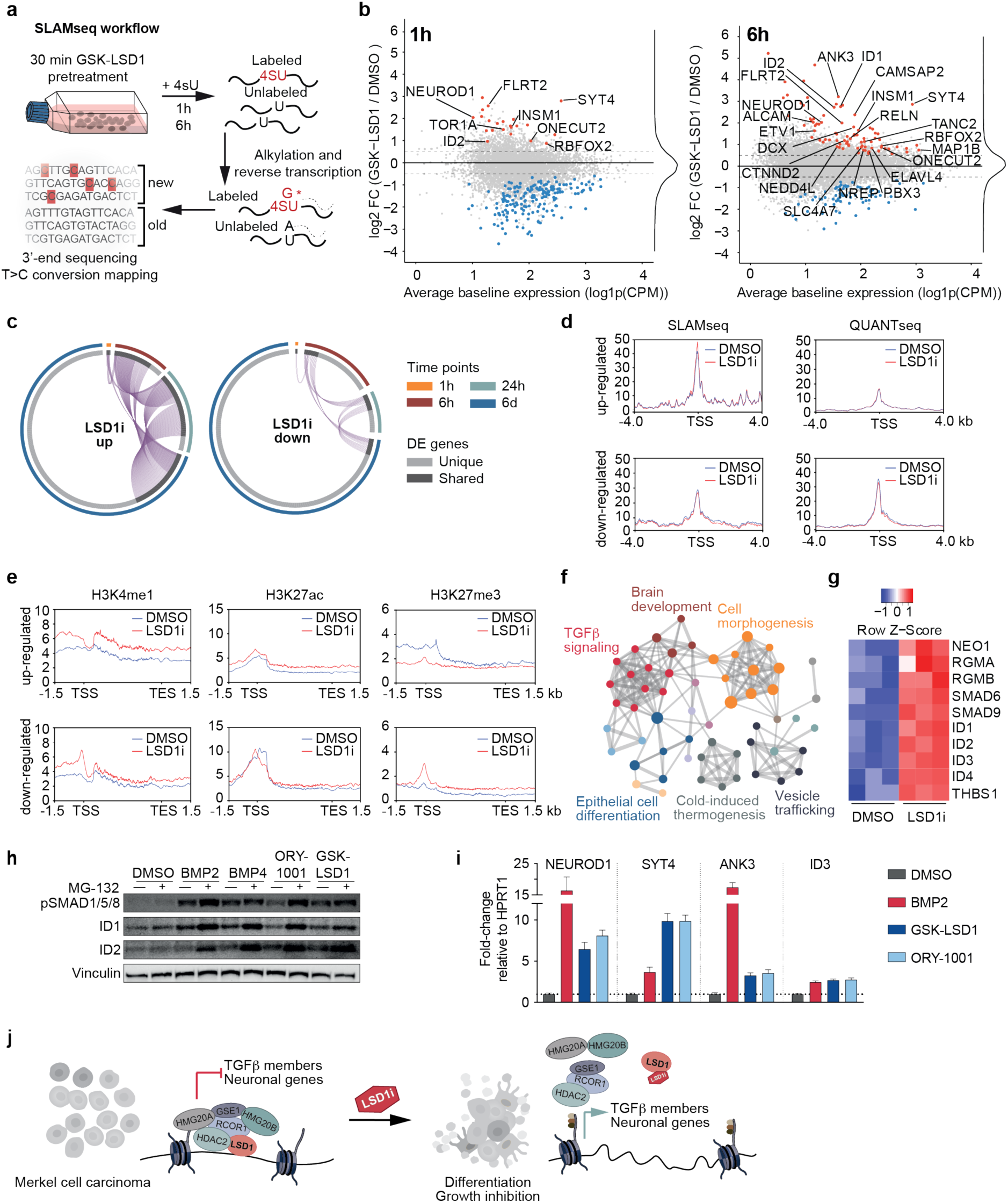
Mapping the direct transcriptional response to LSD1i reveals TGFβ signaling mediated activation of neuronal genes. **a**, Schematic of SLAMseq work-flow. Cells were pretreated for 30 min with LSD1 inhibitor and subsequently labelled with 4-thiouridine (4sU). RNA was extracted and alkylated and subjected to 3’-sequencing. **b**, MA-plot of the mRNA changes detected by SLAMseq after 1h and 6h of 4sU labelling. Significantly up- and downregulated genes (FDR<0.05; abs(log2FC)>0.5) are marked in red and blue, respectively. Genes belonging to neuronal differentiation (GO:0048666) are labelled. FC, fold-change; CMP, counts per million. **c**, Circos plot depicting the transcriptional changes of genes up- (left) or down- (right) regulated upon LSD1i at 1h, 6h, 24 and 6d. **d**, ATAC-seq peak profile of direct SLAMseq targets and differentially expressed (DE) genes at d6 upon LSD1 inhibition or DMSO treatment. TSS, transcription start site. **e**, Metagene plots showing the average CUT&RUN signal for H3K4me1, H3K27ac, and H3K27me3 for all differentially expressed genes upon LSD1i treatment. TSS, transcription start site; TES transcription ed site. **f**, Metascape analysis of upregulated genes for enriched pathway terms in genes identified after 6h of 4SU-labeling. **g**, Heatmap depicting the transcriptional activation of members of TGFβ signaling after 24h treatment with LSD1i. **h**, Western blot probing for protein levels of phospho-SMAD1/5/8, ID1 and ID2 in PeTa cells treated with indicated compounds. Vinculin serves as loading control. **i**, RT-qPCR quantification of neuronal differentiation markers upon 24h BMP2 or LSD1i treatment. **j**, Schematic model of LSD1 mode-of-action in MCC.

**Extended Data Fig. 5.**
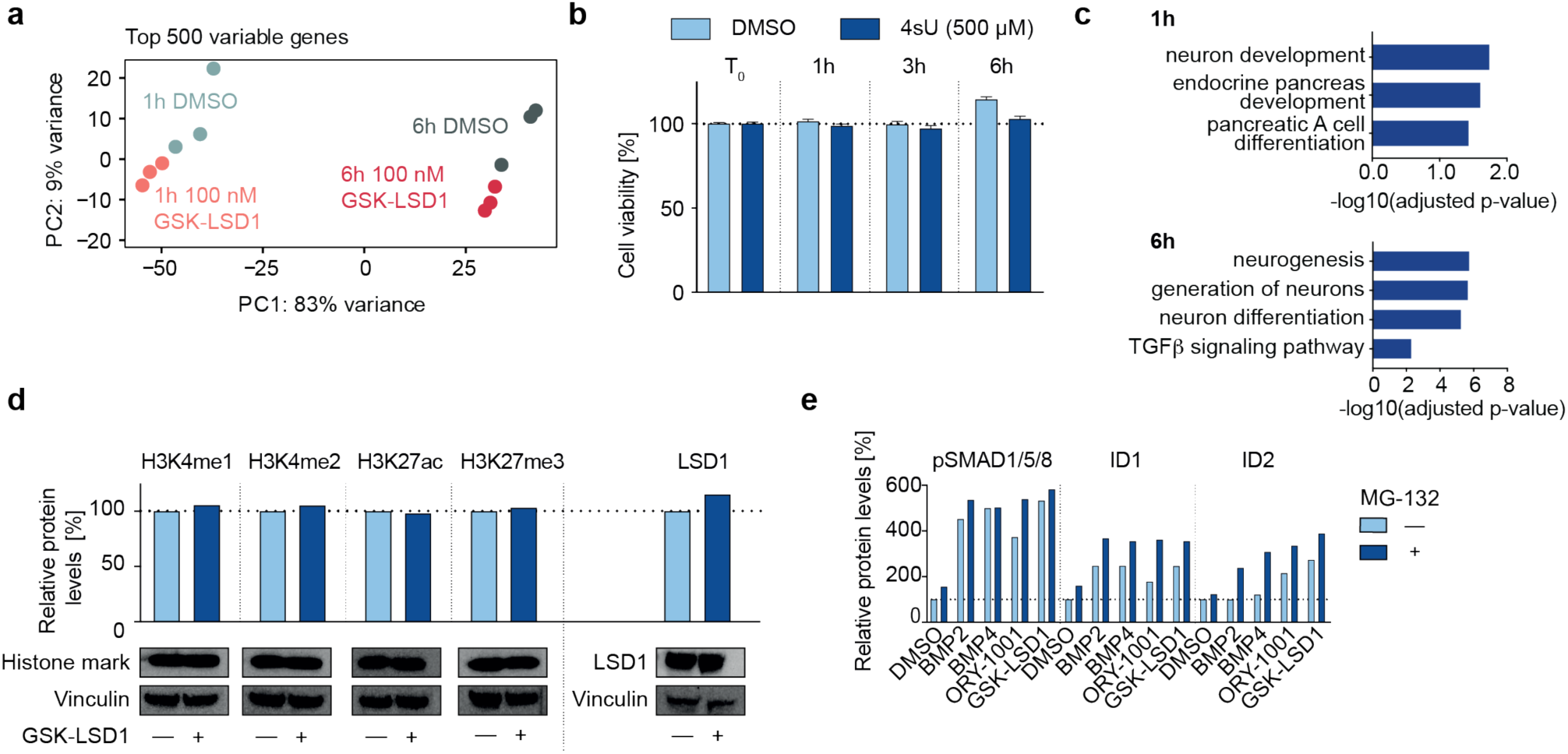
Mapping the direct transcriptional response to LSD1i reveals activation of neuronal genes and TGFβ pathway members. **a**, Principal component analysis (PCA) of the top 500 variable genes in Fig. 5b. PC, principal component. **b**, Cell viability (%) upon 4-thiouridine (4sU) treatment in PeTa cells over time relative to treatment start (T_0_). Data are represented as means ±SD. **c**, Pathway-enrichment analysis for direct transcriptional targets of LSD1 in Fig. 5b. **d**, Westen blot probing for histone marks and LSD1 upon GSK-LSD1 and (bottom) protein level quantification normalized to loading control (vinculin) and relative to DMSO control. **e**, Western blot quantification of protein levels shown in Fig. 5h. Protein levels are expressed as normalized to loading control and relative to corresponding DMSO control.

To assess whether LSD1i treatment alters chromatin accessibility in MCC cells, we performed ATACseq. We found that even long-term LSD1 inhibition (6 d) did not alter chromatin accessibility globally or at the LSD1i response genes inferred from SLAMseq (**Fig. 5d**). LSD1 target genes displayed open chromatin at their transcriptional start sites, independently of LSD1i treatment, suggesting that LSD1 inhibition transcriptionally de-represses these genes in MCC cells. Next, we investigated whether the transcriptional response to LSD1 inhibition changes the histone marks. Immunoblots showed no global changes in histone modifications (**Extended Data Fig. 5d**). We performed

CUT&RUN profiling for the chromatin marks H3K4me1, H3K27ac, and H3K27me3 and assessed the changes in the promoter region and the gene body using metagene plots. At the proximal promoter of genes up-regulated by LSD1i, we found an enrichment for H3K4me1, a modest increase of H3K27ac, and a pronounced decrease of H3K27me3 (**Fig. 5e**).

Pathway enrichment analysis of the immediate transcriptional responses to LSD1i highlighted a prominent cluster of TGFβ pathway members (**Fig. 5f**), including BMP-receptors (NEO1, RGMA, RGMB), the downstream actuator SMAD8/9, and the ID family proteins ID1-4 robustly upregulated after 24h of LSD1i treatment by RNAseq (**Fig. 5g**). Bone morphogenetic protein (BMP) signaling, as part of the TGFβ signaling pathway, regulates cell fate determination and has a well-established role in neuronal differentiation^23^. To determine if BMP-receptor mediated activation of the TGFβ pathway in MCC cells would phenocopy LSD1 inhibition, we treated MCC cells with the ligands BMP2 or BMP4. Indeed, we observed increased levels of phospho-SMAD8/9, ID1 and ID2 in MCC cells treated with BMP2, BMP4 or LSD1i compared to DMSO (**Fig. 5h and Extended Data Fig. 5e**). ID proteins negatively regulate the DNA-binding of basic helix-loop-helix (bHLH) transcription factors by forming heterodimers and contributing to their degradation^24,25^. To investigate the stability of ID1, ID2, and phospho-SMAD8/9, we co-treated MCC cells with the proteasome inhibitor MG-132 and BMP2, BMP4 or LSD1i. Proteasomal inhibition increased the levels of ID1, ID2, and phospho-SMAD8/9, but only in MCC cells treated with BMP2, BMP4 or LSD1i (**Fig. 5h and Extended Data Fig. 5e**). We found that BMP2-mediated activation of TGFβ signaling induces expression of neuronal differentiation markers such as NEUROD1, SYT4 and ANK3 (**Fig. 5i**), which are direct transcriptional targets of LSD1 in MCC. Altogether, our data suggest that transcriptional activation of TGFβ signaling in LSD1i-treated MCCs phenocopies receptor-ligand based TGFβ activation and induces neuronal differentiation.

Our work reveals LSD1 as a previously unappreciated therapeutic target in MCC. We demonstrate that LSD1 inhibitors induce TGFβ signaling and attenuate tumor progression by differentiating MCC to a neuronal lineage, resembling normal Merkel cells (**Fig. 5j**). We show that pharmaceutical inhibition of LSD1 leads to disruption of the LSD1-CoREST complex, including HMG20B. Notably, HMG20B is important for repression of neuronal lineage-specific genes^18,19^. Interestingly, the inhibition of T antigen, the oncogenic driver of the Merkel cell polyomavirus integrated in 80% of MCC, has recently been shown to induce neuronal differentiation when co-cultured with keratinocytes^26^. It is tempting to speculate that T antigen-mediated transformation relies on LSD1 to suppress differentiation towards normal Merkel cell fate and lock MCC cells in a stem-like state. Therapies that target cell fate regulators, instead of aberrantly activated oncogenic drivers e.g. kinases, have not been extensively explored in solid cancers, but demonstrate good clinical activity in leukemias, i.e. all-trans retinoic acid in acute promyelocytic leukemia^27,28^. Here, we provide evidence that therapies targeting cell fate regulators could be powerful in solid cancers too. Importantly, LSD1 inhibitors are currently being tested in clinical trials for other malignancies and could be repurposed for treatment of MCC. The combination of LSD1 inhibitors with therapeutic modalities targeting different cancer hallmarks, such as immune checkpoint inhibitors, could be promising to limit targeted therapy resistance.

## Acknowledgements

We thank all members of the Obenauf lab for experimental support and discussions. We further thanks the core facilities of IMP/IMBA especially the Protein Chemistry for their mass spectrometry expertise, the BioOptics for the FACS sorting, the Molecular Biology Service for reagents and support, the Translational Medicine facility for the animal care and the VBCF Next Generation Sequencing Facility for sequencing. The Epigenetics Probes Collection was supplied by the Structural Genomics Consortium (www.thesgc.org) under an Open Science Trust Agreement. This work was funded by the Starting Grants of the European Research Council (ERC-StG-759590 to A.C.O) and the Vienna Science and Technology fund (#LS16-063 to A.C.O and T.W.). Research at the IMP is generously supported by Boehringer Ingelheim.

## Author contributions

L.L., P.S.J., A.C.O. and T.W. conceived the study, designed the experiments, interpreted the results and wrote the manuscript. A.C.O. and T.W. jointly supervised the study. L.L. and P.S.J. equally contributed to *in vitro* and *in vivo* experiments and computational analysis of the data. T.N. provided computational analysis support. All authors read and approved the manuscript.

## Methods

### DepMap data analysis

The DepMap (https://depmap.org/portal/) RNAi screen dataset (rnai_19Q1, EH2260) was analyzed using the Bioconductor R package “depmap”^29^. Individual or mean dependencies for genes targeted in the epigenetic modifier screen were calculated for the 3 MCC cell lines (MKL-1, MKL-2, PeTa). LSD1 and HMG20B dependency scores across all cancer cell lines were grouped by tissue type as predefined by DepMap. MCC cell lines were removed from the tissue type “SKIN” and grouped as “MCC”. Tissue types with less than three cell lines or for cell lines with less than three dependency scores for a specific gene were removed from the dataset. Violin plots and boxplots were plotted with the R package “ggplot2”^30^.

### Cell culture

The MCC cell lines, MKL-1, MKL-2, MS-1, PeTa, WaGa were cultured as suspension cells in RPMI-1640 supplemented with 10% FBS, 2 mM L-glutamine, 50 U/mL penicillin and 50 mg/mL streptomycin. HDFB and HEK-293T cells were cultured as an adherent monolayer in DMEM-high glucose medium supplemented with 10% FBS, 2 mM L-glutamine, 50 U/mL penicillin and 50 mg/mL streptomycin. All cells were maintained at 37°C and 5% CO2.

### Epigenetic modifier screen

Suspension cells were seeded at a density of 10,000 cells per well; adherent cells were plated at a density of 2,500 cells per well in a 96-well plate. The epigenetic probes collection, obtained from the Structural Genomics Consortium (http://www.thesgc.org) was dissolved in DMSO and compounds were probed at concentrations of 100 µM, 1 µM, 10 nM and DMSO only. Cell viability was determined with the CellTiter-Glo assay (Promega) according to the manufacturer’s instructions on day 6 after seeding. Dose-response curves were generated in quadruplicates. IC50 values were calculated using the GRmetrics R package^31^.

### Dose-response curves

Suspension cells were seeded at a density of 10,000 cells per well; adherent cells were plated at a density of 2,500 cells per well in a 96-well plate. ORY-1001 and GSK-LSD1 were resuspended in DMSO and serially diluted with final concentrations ranging from 0.01 nM to 100 µM. Cells were treated in quadruplicates at indicated doses for 6 days. Cell viability was read out with the CellTiter-Glo assay (Promega) according to the manufacturer’s instructions. IC50 curves were calculated with the software GraphPad PRISM 8 (non-linear regression, log(inhibitor vs response - variable slope, four parameters).

### Photomicrographs

MCC cells (PeTa, WaGa, MS-1) were seeded at a density of 10,000 cells per well in a 96-well plate. Cells were treated with GSK-LSD1 (100 nm) or DMSO. Photomicrographs were taken on day 6. Scale bar was defined with ImageJ.

### Lentivirus production and cell transduction

For lentivirus production, HEK-293T cells were co-transfected with the plasmid of interest, VSVG as envelope plasmid, and the packaging plasmid PAX2 in standard medium with polyethyleneimine (PEI) as previously reported^22^. 24 hours after transfection, the culture medium was changed to 1% FBS medium and viral supernatant was collected after 24 hours and filtered through a 0.4 µm mesh. Transduction of cell lines was performed by spinfection at 800g for 30 min at 32°C with 10 µg/mL polybrene.

### Competition assay

Cells (3e+06) were transduced at a multiplicity of infection (MOI) of ∼1 with shRNAs cloned into a doxycycline-inducible, GFP-expressing and puromycin selectable backbone (T3G-GFP-miRE-PGK-Puro-IRES-rtTA3, LT3GEPIR), resulting in ∼30% GFP-positive cells at day 0 of the experiment. The shRNA expression was induced with 500 ng/mL doxycycline. The relative abundance of transduced cells (GFP-positive cell population versus non-transduced population) was followed over time by flow cytometry (FACS LSR Fortessa). shRNA hairpins targeting Renilla luciferase and RPS15 were used as non-targeting and killing control, targeting a core-essential gene, respectively.

### shRNA cloning

shRNAs were designed with the splashRNA software^32^ and cloned into the backbone by Gibson assembly as previously described^33^. All designed shRNAs, lentiviral backbones and primers used for cloning are listed in **Supplementary Table 3-5**.

### Flow-cytometry

Suspension cells were pelleted at 400g for 5 min, resuspended in Accutase and incubated at 37°C for 5 min. 20,000 single-cell events were acquired per sample on a FACS LSR Fortessa cytometer and GFP-positive fractions were determined with the FlowJo software.

### Immunoblotting

Cells were lysed with RIPA buffer supplemented with cOmplete™ Protease Inhibitor Cocktail (Sigma-Aldrich) and HALT™ phosphatase inhibitor (Thermo Fisher Scientific). Lysates were sonicated and cleared by centrifugation at 14,000g for 10 min at 4°C. Protein concentrations were determined with the BCA Protein Assay kit (Thermo Fisher Scientific) according to the manufacturer’s instructions. Immunoblotting was conducted according to standard protocols. The antibodies used are listed in **Supplementary Table 6**. Protein levels were quantified relative to the loading control with ImageJ and normalized to the loading control.

### RNA extraction

Total RNA isolation from cells was performed with an in-house paramagnetic bead-based purification protocol. Briefly, cells were lysed, incubated with paramagnetic beads and stringently washed to remove proteins. Beads were then treated with DNaseI and purified RNA was eluted in nuclease-free water. Concentration was determined with the spectrophotometer/fluorometer DeNovix DS-11 Fx and with Qubit™ 1X dsDNA HS Assay Kit (Thermo Fisher Scientific).

### qPCR

First-strand synthesis was performed with 1 µg of extracted total RNA with SuperScript™ III Reverse Transcriptase (Invitrogen) according to the manufacturer’s instructions. The qRT-PCR was performed with GoTac® qPCR (Promega) with 10 ng cDNA template on a Biorad CFX384 Real Time Cycler. Data were normalized to the housekeeping gene HPRT1 and displayed as relative to shRenilla. A list of the used primers can be found in **Supplementary Table 5**.

### LSD1-immunoprecipitation

Cells were pre-treated with either 1 µM ORY-1001 or DMSO 24 hours prior to the co-IP. Cells were next washed with ice-cold PBS and lysed on ice with 1x Cell Lysis Buffer (CST). Lysates were cleared by centrifugation at 14,000g for 10 min at 4°C. The lysate concentration was determined with the BCA Protein Assay kit (Thermo Fisher Scientific) according to the manufacturer’s instructions. Protein A sepharose magnetic beads were pre-washed with 1x Cell Lysis Buffer (CST). To reduce nonspecific binding to the beads, the lysates were pre-cleared by incubating with the pre-washed beads for 20 min at RT on a rotator. The pre-cleared lysates were incubated with the anti-LSD1 (CST) primary antibody overnight at 4°C on a rotator to form the immunocomplex. Control samples were incubated with an isotype control antibody. The protein-antibody complexes were then immunoprecipitated for 20 min at RT on a rotator with protein A sepharose magnetic beads which were pre-washed with 1x Cell Lysis Buffer (CST). After incubation, bead-bound immunocomplexes were washed three times with 1x Cell Lysis Buffer (CST) followed by ten PBS washes prior to MS analysis. Confirmation of successful co-IP was conducted by immunoblotting according to standard procedures.

### NanoLC - Mass Spectrometry Analysis

The nano HPLC system used was an UltiMate 3000 RSLC nano system (Thermo Fisher Scientific, Amsterdam, Netherlands) coupled to a Q Exactive HF-X mass spectrometer (Thermo Fisher Scientific, Bremen, Germany), equipped with a Proxeon nanospray source (Thermo Fisher Scientific, Odense, Denmark). Peptides were loaded onto a trap column (Thermo Fisher Scientific, Amsterdam, Netherlands, PepMap C18, 5 mm × 300 µm ID, 5 µm particles, 100 Å pore size) at a flow rate of 25 µL min-1 using 0.1% TFA as mobile phase. After 10 min, the trap column was switched in line with the analytical column (Thermo Fisher Scientific, Amsterdam, Netherlands, PepMap C18, 500 mm × 75 µm ID, 2 µm, 100 Å). Peptides were eluted using a flow rate of 230 nl min-1, and a binary 3h gradient, respectively 225 min.

The gradient starts with the mobile phases: 98% A (water/ formic acid, 99.9/0.1, v/v) and 2% B (water/acetonitrile/formic acid, 19.92/80/0.08, v/v/v), increases to 35%B over the next 180 min, followed by a gradient in 5 min to 90%B, stays there for 5 min and decreases in 2 min back to the gradient 98%A and 2%B for equilibration at 30°C.

The Q Exactive HF-X mass spectrometer was operated in data-dependent mode, using a full scan (m/z range 380-1500, nominal resolution of 60,000, target value 1E6) followed by 10 MS/MS scans of the 10 most abundant ions. MS/MS spectra were acquired using normalized collision energy of 28, isolation width of 1.0 m/z, resolution of 30.000 and the target value was set to 1E5. Precursor ions selected for fragmentation (exclude charge state 1, 7, 8, >8) were put on a dynamic exclusion list for 60 s. Additionally, the minimum AGC target was set to 5E3 and the intensity threshold was calculated to be 4.8E4. The peptide match feature was set to preferred and the exclude isotopes feature was enabled.

### Mass Spectrometry data processing

For peptide identification, the RAW-files were loaded into Proteome Discoverer (version 2.3.0.523, Thermo Scientific). All hereby created MS/MS spectra were searched using MSAmanda v2.3.0.12368, Engine version v2.0.0.12368^34^. For the 1st step search the RAW-files were searched against the swissprot database, taxonomy Homo sapiens (20.341 sequences; 11,361,548 residues), supplemented with common contaminants, using the following search parameters: The peptide mass tolerance was set to ±5 ppm and the fragment mass tolerance to 15ppm. The maximal number of missed cleavages was set to 2, using tryptic enzymatic specificity. The result was filtered to 1 % FDR on protein level using the Percolator algorithm^35^ integrated in the Thermo Proteome Discoverer. A sub-database was generated for further processing.

For the 2nd step, the RAW-files were searched against the created sub-database using the following search parameters: Beta-methylthiolation on cysteine was set as a fixed modification, oxidation on methionine, deamidation of asparagine and glutamine, acetylation on lysine, phosphorylation on serine, threonine, and tyrosine, methylation and di-methylation on lysine and arginine, tri-methylation on lysine, ubiquitination on lysine and biotinylation on lysine were set as variable modifications. Monoisotopic masses were searched within unrestricted protein masses for tryptic enzymatic specificity respectively. The peptide mass tolerance was set to ±5 ppm and the fragment mass tolerance to ±15 ppm. The maximal number of missed cleavages was set to 2. The result was filtered to 1 % FDR on protein level using the Percolator algorithm integrated in Proteome Discoverer. The localization of the post-translational modification sites within the peptides was performed with the tool ptmRS, based on the tool phosphoRS^36^. Peptide areas were quantified using the in-house-developed tool apQuant^37^.

### Metascape analysis

The analysis was performed in single or multi list mode using significantly enriching genes in the respective experiment allowing comparative analysis of datasets across multiple samples^38^.

### HMG20B cloning

cDNA of HMG20B WT was cloned into a lentiviral pTwist backbone encoding for SFFV-V5-cDNA-P2A-mCherry-IRES-Puro-WPRE. At the location of cDNA, we either expressed the wild-type (WT) version of HMG20B or as a control, an empty vector was generated with IRFP670 at the position of cDNA. All vectors were generated by Twist Bioscience and validated by Sanger sequencing.

### HMG20B rescue experiment

PeTa cells were transduced to express either WT HMG20B or empty vector (EV - IRFP670) and selected using 2 µg/ml puromycin. Next, cells were co-transduced with a doxycycline-inducible, shRNA-expressing backbone at a multiplicity of infection (MOI) of ∼1 (T3G-GFP-miRE-PGK-Puro-IRES-rtTA3). shRNA were either targeting endogenous HMG20B or Renilla luciferase. The shRNA expression was induced with 500 ng/mL doxycycline. The abundance of transduced cells was followed by measuring GFP expression over time by flow cytometry (FACS LSR Fortessa).

### SLAMseq

Cells grown at about 50% of the maximum cell density counted on a hemocytometer were pre-treated with a GSK-LSD1 at 100 nM or DMSO for 30 minutes to pre-established full target inhibition. Newly synthesized RNA was labeled by the addition of 500 µM 4-thiouridine (4sU) for 60 or 360 min respectively. Cells were harvested by centrifugation and immediately snap-frozen. RNA extraction was performed using an in-house magnetic beads-based purification protocol as mentioned above. As previously reported^22^, alkylation of total RNA was induced with iodoacetamide (10 mM) for 15 min and RNA was re-purified by ethanol precipitation. As input for generating 3’-end mRNA sequencing (Quantseq) libraries, 500 ng alkylated RNA was used and prepared with a commercially available kit (QuantSeq 3′ mRNA-Seq Library Prep Kit FWD for Illumina and PCR Add-on Kit for Illumina, Lexogen). Sequencing was performed on an Illumina NovaSeq SP platform in 100bp-single-read mode.

### 4-thiouridine toxicity assessment

Cells were treated with 500 µM 4-thiouridine (4sU) or with DMSO. Cell viability was read out with the CellTiter-Glo assay (Promega) according to the manufacturer’s instructions at treatment start (T0), after 1 hour, 3 hours and 6 hours. Cell viability at each time point was calculated relative to the respective measure at T0.

### SLAMseq data analysis

Data analysis was performed as previously reported^22^. Briefly, gene and 3’ UTR annotations for hg38 were obtained from UCSC table browser. Adapters were trimmed from raw reads using cutadapt through the trim_galore (v0.3.7) wrapper tool with adapter overlaps set to 3bp for trimming. Trimmed reads were further processed using SlamDunk^39^ using the (slamdunk all) method. Reads were filtered for having ≥ 2 T>C conversions. Differentially expressed genes were calculated using DESeq2 (v1.18.140). Pathway analysis was performed using gProfiler^41^ and Metascape^38^.

### QUANTseq

Cells were treated for 24h or 6 days with an LSD1 inhibitor (GSK-LSD1, 100 nM) or DMSO. At harvest time points, cells were spun down, washed in PBS and cell pellets were snap-frozen. RNA extraction was performed using an in-house magnetic beads-based purification protocol mentioned above. As input for generating 3’-end mRNA sequencing (Quantseq) libraries, 500 ng RNA was prepared with a commercially available kit according to the manufacturer’s instruction (QuantSeq 3′ mRNA-Seq Library Prep Kit FWD for Illumina and PCR Add-on Kit for Illumina, Lexogen). Sequencing was performed on an Illumina HiSeq2500 in 50bp single-read mode.

### QUANTseq data analysis

Analysis of QUANTseq data was performed with an in-house pipeline, briefly: Adapter and polyA sequences were clipped (bbdk, 38.06, https://sourceforge.net/projects/bbmap/), abundant sequences were removed (bbmap, 38.06; https://sourceforge.net/projects/bbmap/) and cleaned reads were aligned against the genome (hg38) with (STAR, 2.6.0c^42^). Raw reads were mapped to 3’ UTR-annotations of the same gene and collapsed to gene level by Entrez Gene ID with featureCounts (v1.6.2). Differentially expressed genes were calculated using DESeq2 (v1.18.1^40^). Pathway analysis was performed using gProfiler^41^ and Metascape^38^.

### Gene set enrichment analysis (GSEA)

Gene set enrichment analysis^43^ was performed using the GSEA tool (4.0.3) with a normalized gene expression table and standard parameters for enrichment testing.

### Motif analysis

Gene-based motif analysis was performed using Homer (4.11)^44^ with the findMotifs.pl script and default settings. The top 5-enriched motifs enriched in genes up- or down-regulated were plotted.

### CUT&RUN

CUT&RUN was performed according to the high-calcium/low-salt digestion protocol described previously^45^. Briefly, cells were harvested, washed and bound to activated Concanavalin A-coated magnetic beads and permeabilized (20 mM HEPES, pH 7.5, 150 mM NaCl, 0.5 mM spermidine, Roche cOmplete™ and 0.05 % Digitonin). The bead-cell complex was incubated overnight with the respective antibody at 4°C. Cells were washed three times and resuspended in 100 µl pAG/MNase and incubated for 1h at RT. Antibodies used are listed in **Supplementary Table 6**.

### CUT&RUN data analysis

CUT&RUN data was analyzed utilizing the nf-core^46^ ChIPSeq pipeline^47^ using the human genome hg38 and calling peaks in broad-peak mode. Aligned reads of the individual replicates were merged using samtools v1.9^48^ and RPGC-normalized tracks were calculated and plotted as composite density plots using deeptools v3.1.2^49^. Enhancers and super-enhancers on the CUT&RUN data were called using the ROSE^50^ tool (v0.1) excluding TSS within 2.5kb.

### ATACseq

ATACseq was performed according to the Omni-ATACseq protocol^51^. Briefly, 100,000 cells were pelleted, and resuspended in resuspending buffer (10 mM Tris-HCl, 10 mM NaCl, 3 mM MgCl2, 0.1% Nonidet P40, 0.1% Tween-20 and 0.01% digitonin). After nuclei isolation cells were resuspended in TD buffer with in-house purified Tn5 transposase. The transposition reaction was performed at 37°C for 30 min. DNA fragments were purified from the reaction using a Zymo DNA Clean & Concentrator-5 kit (in-house) and amplified with NEBNext High Fidelity PCR Mix (NEB). Library quality was assessed using a Fragment Analyzer system (Agilent). ATACseq libraries were quantified using a KAPA library quantification kit and sequenced on an Illumina NextSeq550 instrument in 75 bp paired-end mode.

### ATACseq data analysis

ATACseq data was analyzed using the nf-core/ATACseq pipeline^52^ aligning against the human genome hg38 and calling peaks with MACS2 in narrow- and broad-peak mode. Aligned reads of the individual replicates were merged using samtools v1.9^48^ and RPGC-normalized tracks were calculated and plotted as composite density plots using deeptools v3.1.2^49^.

### TGFβ signaling pathway induction

Cells were pretreated with either active BMP2 recombinant protein (10 ng/mL), active BMP4 recombinant protein (10 ng/mL), ORY-1001 (1 µM), GSK-LSD1 (1 µM) or DMSO for 24 hours. Prior to protein and RNA harvest, each condition was split and was either co-treated with the proteasome inhibitor MG-132 (1 µM) or DMSO for 2 hours. Cells were pelleted, washed with PBS and lysed with RIPA buffer as described above. Immunoblotting and RT-qPCR were conducted according to standard procedures.

### PARP1 cleavage

Cells were treated ORY-1001 (10 nM or 100 nM), with the positive control for apoptosis etoposide (10 µM) or DMSO for 24 hours. Cells were afterwards harvested, washed with PBS and lysed with RIPA buffer as described above. Immunoblotting was conducted according to standard procedures.

### CaspaseGlo readout

Cells were treated either with ORY-1001 (10 nM), GSK-LSD1 (10 nM), or DMSO for 6 days. As a positive control for apoptosis, cells were treated with etoposide (10 µM) for 24 hours. Apoptosis was assessed with Caspase-Glo 3/7 assay (Promega) according to the manufacturer’s instruction.

### KI-67 staining

Cells were treated either with ORY-1001 (1 µM), GSK-LSD1 (1 µM) or DMSO for 3 days. Cells were harvested for 5 min at 400g at RT and stained with Pacific Blue™ anti-human Ki-67 Antibody (Biolegend) according to eBioscience(™) Transcription Factor Staining Buffer Set. 20,000 single-cell events were acquired per sample on a FACS LSR Fortessa cytometer and Pacific Blue(™)-positive fractions were determined with the FlowJo software.

### *In vivo* tumor growth control experiment

PeTa cells (2e+06) mixed with Matrigel were injected subcutaneously into the left flank of five-week-old NSG mice. Prior to tumor cell injection, the mice were injected intraperitoneally with analgesic ketamine (100 mg/kg), xylazine (10 mg/kg) and acepromazine (3 mg/kg). Animal weight was frequently monitored. Tumor size was monitored by tumor measurement with an electronic caliper. Measured tumor size was determined by the formula [x (mm)^2^ * y (mm)]/2 = volume (mm^3^), where x refers to the shortest measurement and y the longest. Once tumor size reached 50 mm^3^ (day 16 post-tumor injection), mice were either treated with GSK-LSD1 (1 mg/kg) or vehicle (0.9% saline) by intraperitoneal (IP) injection. Mice were treated during five consecutive days followed by two days off-treatment (5 days ON / 2 days OFF) repeatedly. Mice were sacrificed by cervical dislocation when either total tumor volume was larger than 1500 mm^3^ or when one of the measured lengths was larger than 1500 mm. All experiments using animals were performed in accordance with our protocol approved by the Austrian Ministry (BMBWF-66.015/0009-V/ 3b/2019 or GZ: 340118/2017/25). Kaplan-Meier survival curves were plotted with the software GraphPad PRISM 8 and significance determined by log-rank Mantel-Cox test. The significance of the grouped xenograft growth was determined by two-way ANOVA.

### *In vivo* micrometastatic settings modeling experiment

PeTa cells (2e+06) mixed with Matrigel were injected subcutaneously into the left flank of five-week-old NSG mice. Prior tumor injection, the mice were injected intraperitoneally with analgesic ketamine (100 mg/kg), xylazine (10 mg/kg) and acepromazine (3 mg/kg). Animal weight was frequently monitored. Tumor size was monitored by tumor measurement with an electronic caliper. Measured tumor size was determined by the formula [x (mm)^2^ * y (mm)]/2 = volume (mm^3^), where x refers to the shortest measurement and y the longest. Mice were either treated with GSK-LSD1 (1 mg/kg), ORY-1001 (0.03 mg/kg) or vehicle (0.9% saline) by intraperitoneal (IP) injection from day one after tumor injection on. Mice were treated during five consecutive days followed by two days off-treatment (5 days ON / 2 days OFF) repeatedly. Mice were sacrificed by cervical dislocation when either total tumor volume was larger than 1500 mm^3^ or when one of the measured lengths was larger than 1500 mm. All experiments using animals were performed in accordance with our protocol approved by the Austrian Ministry (BMBWF-66.015/0009-V/3b/2019 or GZ: 340118/2017/25).. Survival curves were plotted with the software GraphPad PRISM 8 and significance determined by log rank Mantel-Cox test. The significance of the grouped xenograft growth was determined by two-way Anova.

### Merkel cell isolation and sequencing

Dorsal skin was removed from euthanized 4-week-old C57BL/6 mice and dissociated with the Epidermis Dissociation Kit (Miltenyi Biotec, Cat. 130-095-928) according to the manufacturer’s instructions. Briefly, the dorsal skin patches were first incubated with enzyme G to dissociate the epidermis from the dermis. Next, the epidermis was dissociated into a single cell suspension with the gentleMACS™ Dissociator and enzymes P and A. The Merkel cell surface marker NCAM (neural cell adhesion molecule, CD56) was used for the enrichment of a Merkel cell population. The NCAM^+^ Merkel cells in our cell suspension were bound by anti-PSA-NCAM microbeads (Miltenyi Biotec, Cat. 130-097-859) and extracted with the MACS® Column system. Then the Merkel cell-enriched population was centrifuged for 5 min at 400g and resuspended in FACS buffer (PBS supplemented with 0.5% bovine serum albumin (BSA) and 2 mM EDTA). Merkel cells were stained with anti-PSA-NCAM-PE antibody (Miltenyi Biotec, Cat. 130-117-394) or with mouse IgM-PE isotype control (Miltenyi Biotec, Cat. 130-120-070) according to manufacturer’s instructions, for 10 min at 4°C. Cells were washed with FACS buffer and centrifuged for 5 min at 4°C. Next, to exclude dead cells, Merkel cells were resuspended in FACS buffer and stained with DAPI (1 µg/ml) for 30 min at 4°C. NCAM-PE^+^ DAPI-Merkel cells, as well as bulk epidermis cells (DAPI^-^), were FACS-sorted in triplicates, each with 500 events in triplicates in a 96-well plate. The gating strategy for the FACS sorting is depicted in Extended data Fig. 6. Smart-seq2^53^ libraries were prepared and sequenced on an Illumina HiSeqV4 in 50bp single-read mode.

**Extended Data Fig. 6.**
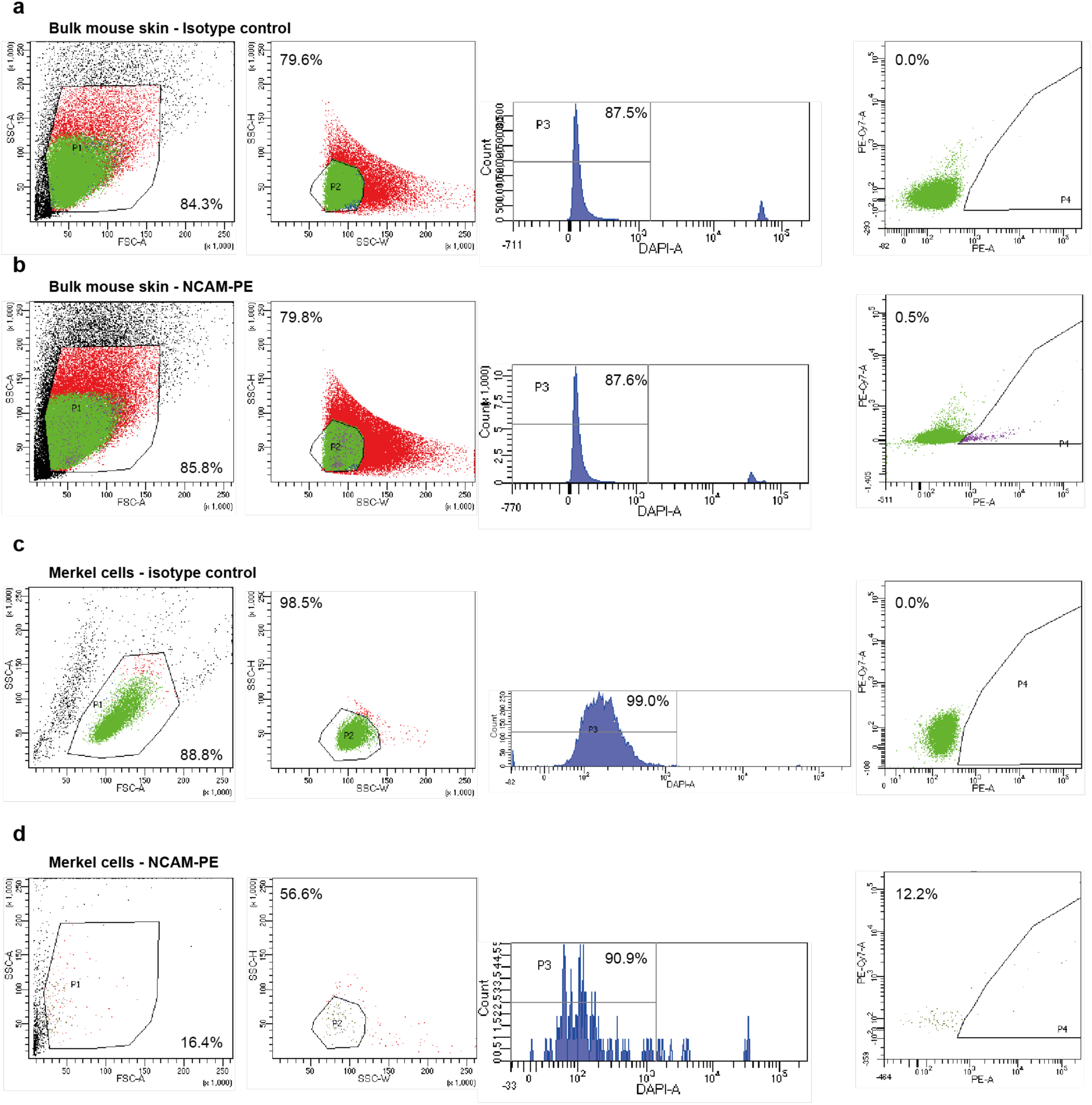
Merkel cell FACS sorting gating strategy for SMART-seq2 sequencing. **a**, Bulk mouse skin labeled with IgM-PE mouse isotype control. **b**, Bulk mouse skin labeled with NCAM-PE. DAPI^-^ cells were sorted. **c**, Merkel cells labeled with IgM-PE mouse isotype control. **d**, Merkel cells labeled with NCAM-PE. DAPI^-^/NCAM^+^ cells were sorted. Each plot shows the subpopulation gated for in the preceding plot to the left.

### SMARTseq analysis

Analysis of SMARTseq data was performed with an in-house pipeline, briefly: Adapter and polyA sequences were clipped (bbdk, 38.06, https://sourceforge.net/projects/bbmap/) abundant sequences were removed (bbmap, 38.06, https://sourceforge.net/projects/bbmap/) and cleaned reads were aligned against the genome (GRCm38) with (STAR, 2.6.0c^42^). Aligned reads were counted with featureCounts (v1.6.2). Differentially expressed genes were calculated using DESeq2 (v1.18.1^40^).

### Data availability

All datasets generated in this study are available from the corresponding authors upon reasonable request.

